# SARS-CoV-2 variants of concern infect the respiratory tract and induce inflammatory response in wild-type laboratory mice

**DOI:** 10.1101/2021.09.29.462373

**Authors:** Shannon Stone, Hussin A. Rothan, Janhavi P. Natekar, Pratima Kumari, Shaligram Sharma, Heather Pathak, Komal Arora, Tabassum T. Auroni, Mukesh Kumar

## Abstract

The emergence of new severe acute respiratory syndrome coronavirus-2 (SARS-CoV-2) variants of concern poses a major threat to the public health due to possible enhanced virulence, transmissibility and immune escape. These variants may also adapt to new hosts in part through mutations in the spike protein. In this study, we evaluated the infectivity and pathogenicity of SARS-CoV-2 variants of concern in wild-type C57BL/6 mice. Six-week-old mice were inoculated intranasally with a representative virus from the original B.1 lineage or emerging B.1.1.7 and B.1.351 lineages. We also infected a group of mice with a mouse-adapted SARS-CoV-2 (MA10). Viral load and mRNA levels of multiple cytokines and chemokines were analyzed in the lung tissues on day 3 after infection. Our data show that unlike the B.1 virus, the B.1.1.7 and B.1.351 viruses are capable of infecting C57BL/6 mice and replicating at high concentrations in the lungs. The B.1.351 virus replicated to higher titers in the lungs compared to the B.1.1.7 and MA10 viruses. The levels of cytokines (IL-6, TNF-α, IL-1β) and chemokine (CCL2) were upregulated in response to the B.1.1.7 and B.1.351 infection in the lungs. Overall, these data indicate a greater potential for infectivity and adaptation to new hosts by emerging SARS-CoV-2 variants.

## 1. Introduction

Coronaviruses are a family of positive-sense single strand RNA viruses whose large genomes and propensity for mutation have resulted in a diversity of strains that are capable of adaptation to new hosts. COVID-19, the disease caused by the new beta coronavirus, SARS-CoV-2, has caused significant human and economic burden **[1-3]**. Few therapies are available to treat COVID-19 disease in humans and the rapid evolution of SARS-CoV-2 variants threatens to diminish their efficacy **[2,4]**. The lineage B.1.1.7, first identified in the United Kingdom, and lineage B.1.351, first described in South Africa, have been termed variants of concern because of the greater risk they pose due to possible enhanced transmissibility, disease severity and immune escape **[4-7]**. These variants may also adapt to new hosts in part through mutations on the receptor binding domain (RBD) of the spike protein **[6,7]**.

SARS-CoV-2 infection begins with the viral particles binding to the receptors on the host cell surface. The RBD of the spike protein binds to angiotensin-converting enzyme 2 (ACE-2) present on the host cellular surfaces **[3,8]**. The RBD of the spike protein from the SARS-CoV-2 strain (Wuhan strain, lineage B.1) that started the pandemic does not efficiently bind mouse ACE-2, and therefore wild-type laboratory mice are not susceptible to infection with lineage B.1 virus **[8-12]**. Mouse-adapted SARS-CoV-2 variant (MA10) with binding affinity to mouse ACE-2 has been obtained after sequential passaging of virus in mouse lung tissues **[11]**. Infection of wild-type BALB/c mice with MA10 virus resulted in replication in both upper and lower airways **[11,13]**. MA10 virus has several mutations, including multiple mutations in the spike protein compared to the Wuhan reference sequence. These mutations are also present in the B.1.1.7 and B.1.351 lineages **[6,7,11,14,15]**. Since the spike protein is necessary for direct interaction with the host receptor, mutations in the spike protein can affect SARS-CoV-2 infection efficiency depending on the host. However, the infectivity and pathogenicity of these emerging variants in mice have not yet been determined.

In this study, we evaluated the replication and pathogenicity of the original B.1 lineage and emerging SARS-CoV-2 lineages, B.1.1.7 and B.1.351, in wild-type C57BL/6 mice. We also used a mouse-adapted SARS-CoV-2 variant (MA10) that causes disease in the wild-type mice **[11]**. Our data show that the B.1.1.7 and B.1.351 viruses are capable of infecting wild-type C57BL/6 mice and replicating at high concentrations in the lungs. The B.1.351 virus replicated to higher titers in the lungs compared to the B.1.1.7 and MA10 viruses. We found that infection with the B.1.351, B.1.1.7 and MA10 viruses trigger an inflammatory response in the lungs characterized by upregulation of inflammatory cytokines and chemokines.

## 2. Materials and Methods

### 2.1. Animal infection experiments

C57BL/6 mice were purchased from the Jackson Laboratory (Bar Harbor, ME). All the animal experiments were conducted in a certified Animal Biosafety Level 3 (ABSL-3) laboratory at Georgia State University (GSU). The protocol was approved by the GSU Institutional Animal Care and Use Committee (Protocol number A20044). Six-week-old C57BL/6 mice were inoculated intranasally with PBS (mock) or 10^5^ plaque-forming units (PFU) of SARS-CoV-2 as described previously [16]. We used B.1 Wuhan virus (BEI# NR-52281), B.1.1.7 virus (BEI# NR-54000), B.1.351 virus (BEI# NR-54008) and MA10 virus (BEI# NR-55329). Roughly equal numbers of male and female mice were used. Animals were weighed and their appetite, activity, breathing and neurological signs assessed twice daily. In independent experiments, mice were inoculated with PBS (Mock) or SARS-CoV-2 intranasally, and on day 3 after infection, animals were anesthetized using isoflurane and perfused with cold PBS. Lungs were collected, and flash-frozen in 2-methylbutane (Sigma, St. Louis, MO, USA) for further analysis as described below **[16,17]**.

### 2.2. Quantification of the virus load

The virus titers were analyzed in the lungs by plaque assay and quantitative real-time PCR (qRT-PCR). Briefly, frozen tissues were weighed and homogenized in a bullet blender (Next Advance, Averill Park, NY, USA) using stainless steel beads. Virus titers in tissue homogenates were measured by plaque assay using Vero cells **[16,17]**. Quantitative RT-PCR was used to measure viral RNA levels with primers and probes specific for the SARS-CoV-2 N gene as described previously **[18]**. Viral genome copies were determined by comparison to a standard curve generated using a known amount of RNA extracted from previously titrated SARS-CoV-2 samples **[18]**. Frozen tissues harvested from mock and infected animals were weighed and lysed in RLT buffer (Qiagen), and RNA was extracted using a Qiagen RNeasy Mini kit (Qiagen, Germantown, MD, USA) **[17,19]**. Total RNA extracted from the tissues was quantified and normalized, and viral RNA levels per !g of total RNA were calculated.

### 2.3. Analysis of cytokines and chemokines

Total RNA was extracted from lungs using a Qiagen RNeasy Mini kit (Qiagen, Germantown, MD, USA). cDNA samples were prepared using an iScript™ cDNA Synthesis Kit (Bio-Rad). The expression levels of multiple host genes were determined using qRT-PCR, and the fold change in infected lungs compared to mock-infected controls was calculated after normalizing each sample to the level of the endogenous GAPDH gene mRNA **[16,17,19]**. The primer sequences used for qRT-PCR are listed in Table 1.

**Table 1.**
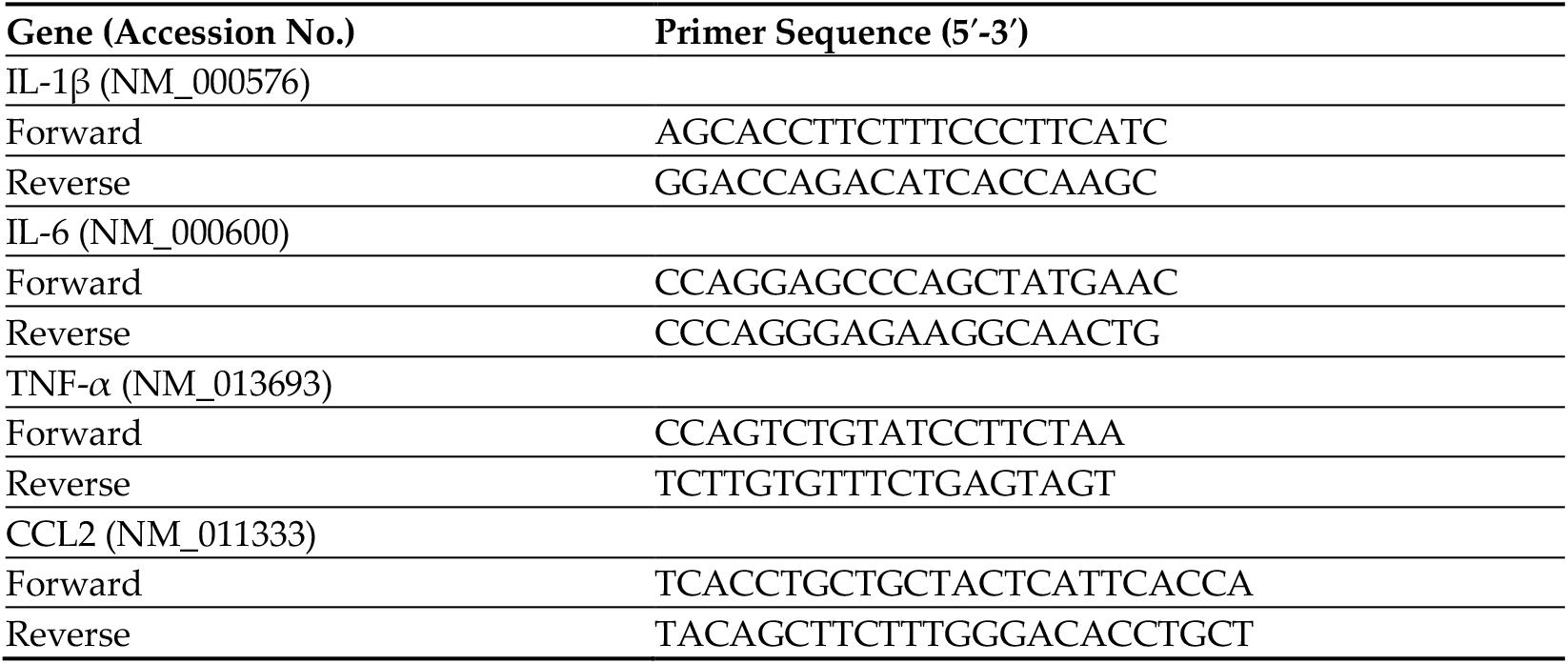
Primer sequences used for qRT-PCR.

### 2.4. Immunoblot analysis

Total cellular protein was extracted from mouse lungs and separated by SDS-PAGE, transferred onto PVDF membranes and incubated overnight with polyclonal antibody against IL-6 **[16,17]**. Membranes were stripped and re-probed with antibody against β-actin. Following incubation with secondary antibodies conjugated with IRDye 800 and IRDye 680 (Li-Cor Biosciences), the membranes were scanned using the Odyssey infrared imager (Li-Cor Biosciences) **[16,17]**.

### 2.5. Statistical Analysis

Mann-Whitney test and unpaired student t-test using GraphPad Prism 5.0 were used to calculate p values of the difference between viral titers and immune responses, respectively. Differences of p <0.05 were considered significant.

## 3. Results

### 3.1. B.1.1.7 and B.1.351 variants replicate in the lungs of C57BL/6 mice

We inoculated six-week-old C57BL/6 mice with 105 PFU of SARS-CoV-2 or PBS (mock) via the intranasal route. We used a representative virus from the original B.1 lineage or emerging B.1.1.7 and B.1.351 lineages. We also infected a group of mice with a mouse-adapted SARS-CoV-2 variant (MA10) **[11]**. Mice were monitored daily for weight loss. Mice infected with B.1 virus exhibited no changes in the body weight. However, B.1.1.7-, B.1.351- and MA10-infected mice had approximately 10% body weight loss at days 2-4 after infection (Figure 1). To evaluate virus replication in the lungs, groups of 5-8 mice were euthanized at 3 days after infection and lungs were collected. Viral infectivity titers in the lungs were measured by plaque assay. Virus was not detected in the mock-infected lungs. High levels of infectious virus were detected in the lungs of all the MA10-infected mice (105 PFU/gram) (Figure 2A). By contrast, little if any infectious virus was detected in the lungs of the B.1-infected mice suggesting limited virus replication. High levels of infectious virus were detected in the lungs of both B.1.1.7- (104 PFU/gram) and B.1.351-infected mice (106 PFU/gram). The viral load was significantly lower for the B.1.1.7 compared to the B.1.351 virus. Virus titers in the lungs of the B.1.351-infected mice were higher than the MA10-infected mice, but the difference was not statistically significant. Intracellular SARS-CoV-2 RNA levels assessed by qRT-PCR followed a similar pattern. The viral RNA levels in the B.1.1.7-, B.1.351- and MA10-infected groups were significantly higher than those in the B.1-infected group (Figure 2B). These data indicate that MA10, B.1.1.7 and B.1.351 viruses replicated efficiently in the lungs of the infected animals, and the B.1.351 virus replicated to high titers in the lungs of mice compared to other groups.

**Figure 1.**
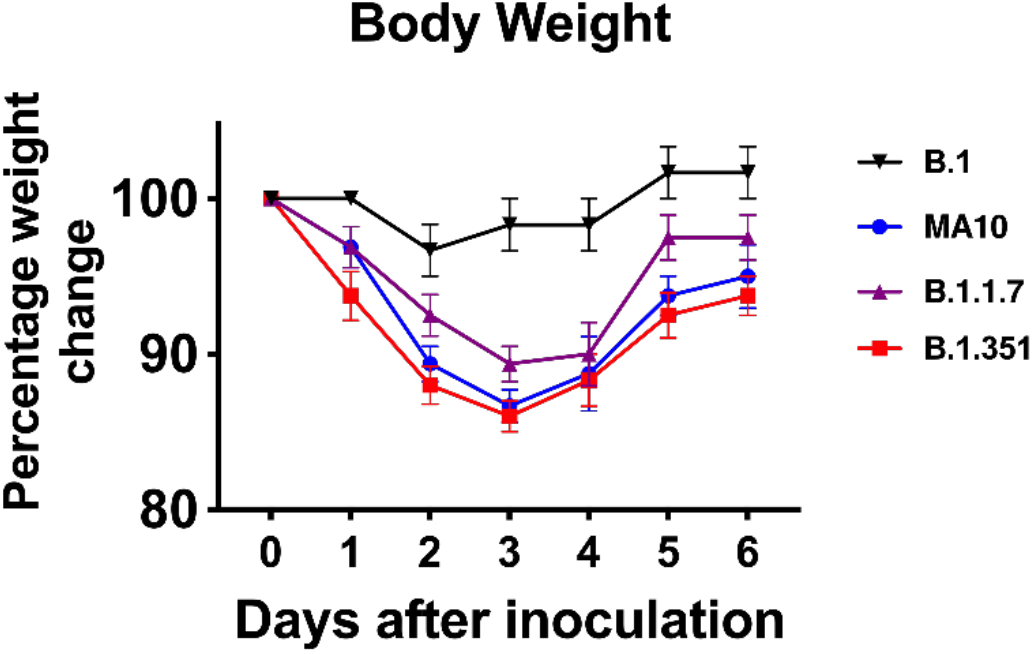
Analysis of body weight in C57BL/6 mice following SARS-CoV-2 infection. Six-week-old C57BL/6 mice were inoculated intranasally with 10^5^ plaque-forming units (PFU) of SARS-CoV-2 variants (n = 8-10 mice per group). Percentage of initial weight for SARS-CoV-2-infected mice over 6 days. Values are the mean ± SEM.

**Figure 2.**
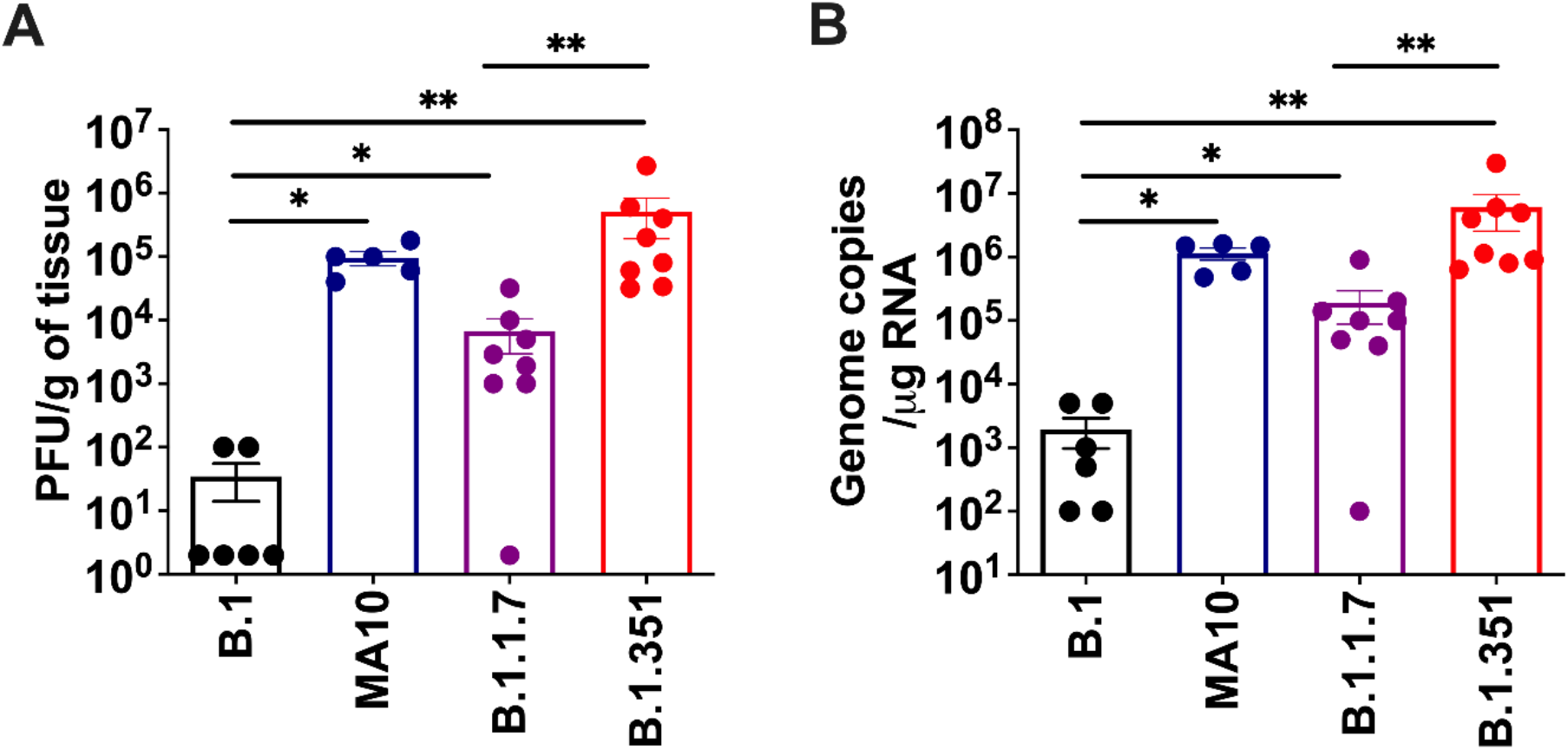
Replication of SARS-CoV-2 variants in the lungs. Six-week-old C57BL/6 mice were inoculated intranasally with PBS (mock) or 10^5^ plaque-forming units (PFU) of SARS-CoV-2 variants. Groups of 5-8 mice were euthanized at 3 days after infection and lungs were collected. Virus titers were analyzed in the lungs by (A) plaque assay and (B) qRT-PCR. The data are expressed as PFU/g of tissue or genome copies/μg of RNA. Each data point represents an individual mouse. *,p < 0.05; **, p< 0.001.

### 3.2. B.1.1.7 and B.1.351 variants induce an inflammatory response in the lungs

The excessive inflammatory host response to SARS-CoV-2 infection contributes to pulmonary pathology and the development of respiratory distress in some COVID-19 patient **[20,21]**. To compare the host responses in SARS-CoV-2-infected mice, we investigated changes in mRNA levels of IL-6, TNF-α, IL-1β and CCL2 in the lungs of mice infected with SARS-CoV-2. Gene expression changes in the lungs of infected mice compared to mock-infected controls were analyzed by qRT-PCR. No significant increase in the mRNAs of the cytokines or chemokines tested was observed in the B.1-infected mice. MA10 infection resulted in a 20-fold increase in IL-6 and CCL2 mRNA expression (Figure 3). IL-1β and TNF-α mRNA levels increased by 4-6-fold in the lungs of the MA10-infected mice. The B.1.351 infection resulted in a >25-fold increase in IL-6 and CCL2 mRNA expression. Similarly, TNF-α mRNA levels increased by 10-fold in the B.1.351-infected mice. There was also a 6-fold increase in IL-1β mRNA in the B.1.351-infected mice. In the B.1.1.7-infected mice, the mRNA levels of the IL-6 and CCL2 were elevated by 10-fold. Although the expression of IL-1β mRNA did not increase, a modest three-fold increase in the TNF-α mRNA level was observed in the lungs of the B.1.1.7-infected mice. Inflammatory response observed in the B.1.351-infected group was significantly higher than in the B.1.1.7- and B.1-infected group. We also measured protein levels of IL-6 in mock- and B.1.351-infected mice at day 3 after infection. Immunoblotting data showed an increase in the protein levels of IL-6 in the B.1.351-infected lungs compared to the mock-infected controls (Figure 4). These results indicate that infection with B.1.351 and B.1.1.7 variants upregulates the expression of inflammatory genes in the lungs.

**Figure 3.**
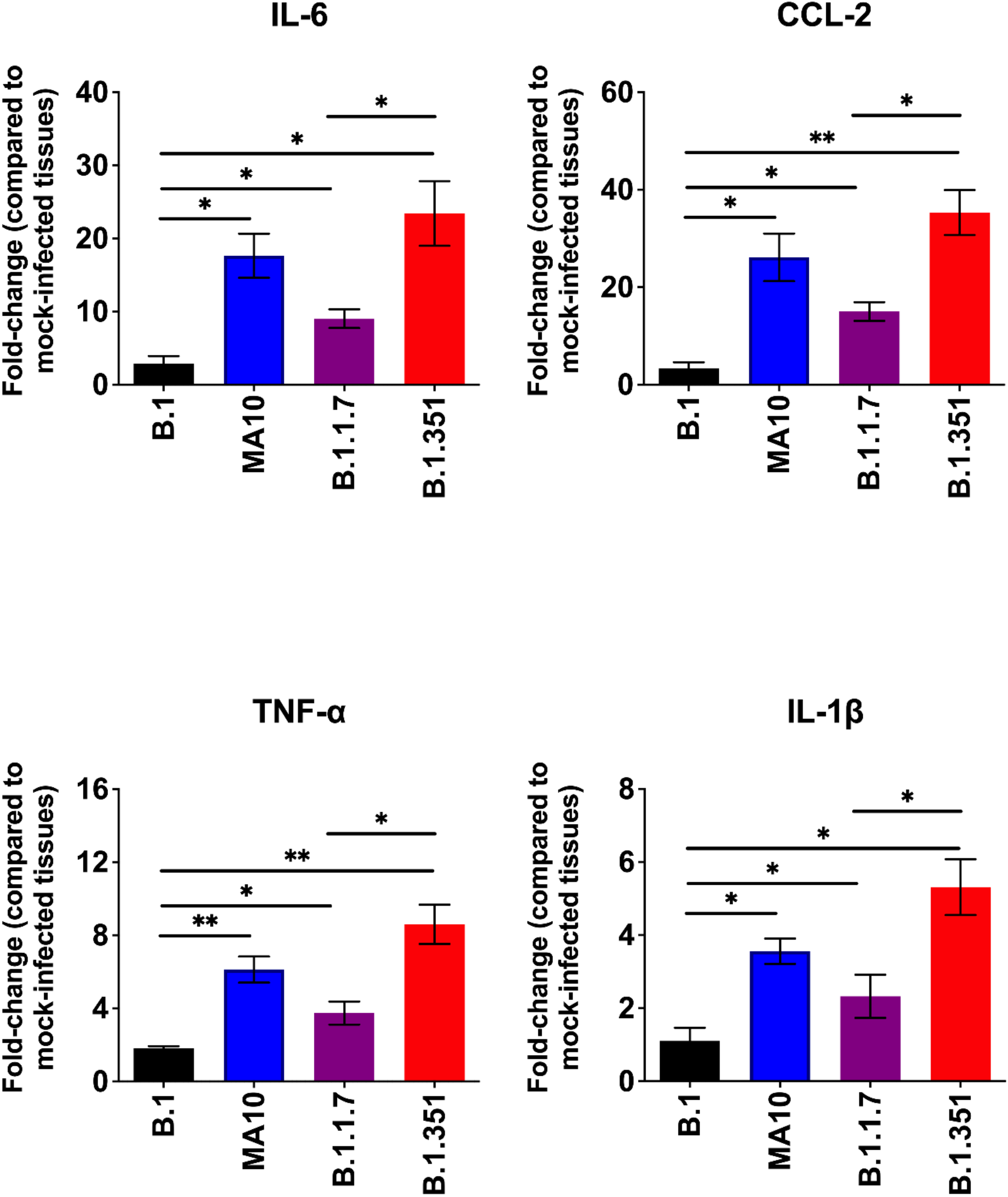
Analysis of inflammatory cytokine and chemokine levels in the lungs of SARS-CoV-2-infected mice. The mRNA levels of IL-6, CCL2, TNF-α and IL-1β genes were determined by qRT-PCR. The fold change in the infected tissues compared to the corresponding mock-infected controls was calculated after normalizing individual samples to GAPDH levels. Values are the mean ± SEM (n = 5-8 mice per group). *, p< 0.05; **, p< 0.001.

**Figure 4.**
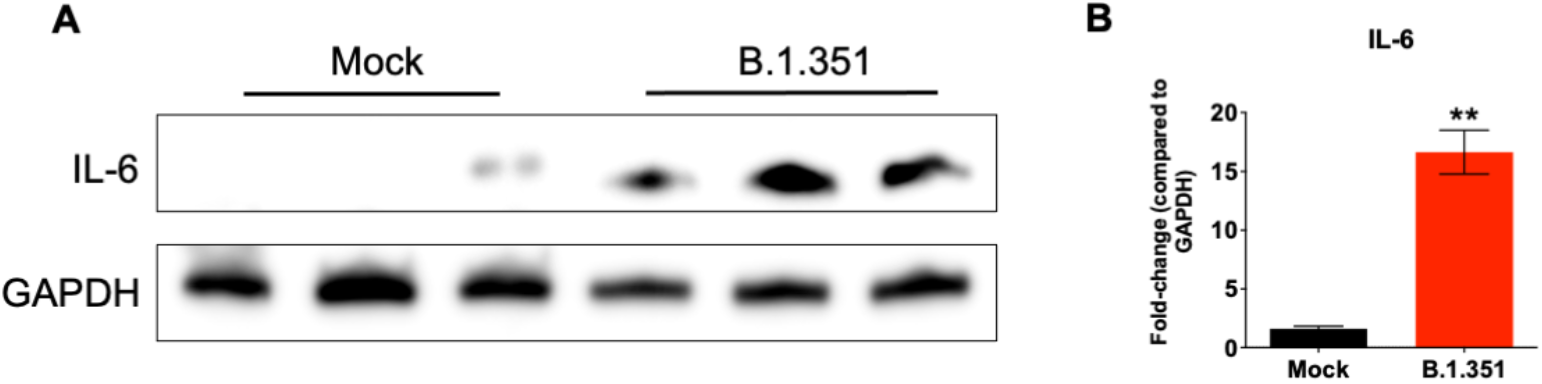
Analysis of protein levels of IL-6 in the lungs of B.1.351-infected mice. (A) Protein extracted from mock- and B.1.351-infected lung tissues (day 3 after infection) were immunoblotted with antibodies against IL-6 or β-actin. Data are representative of five animals per group. (B) Quantitative analysis of Western blot results represented as fold-change compared to GAPDH. **p,0.001 as compared to mock controls (n = 5 mice per group).

## 4. Discussion

This study demonstrates that wild-type laboratory mice are susceptible to infection with the emerging SARS-CoV-2 variants. While the B.1 virus was unable to infect C57BL/6 mice, the B.1.351 and B.1.1.7 viruses efficiently infected C57BL/6 mice. Enhanced virus replication in the B.1.351- and B.1.1.7-infected mice was accompanied by elevated cytokine and chemokine levels in the lungs.

Mice are a useful small animal model for the evaluation of vaccines, immunotherapies and antiviral drugs **[8,12]**. Because the initial SARS-CoV-2 strains do not utilize murine ACE-2 as a receptor, wild-type mice are not susceptible to SARS-CoV-2 infection **[8-10,12]**. Transgenic mice that express human ACE-2 often develop severe and fatal disease upon intranasal inoculation of virus **[16,22]**. The initially available SARS-CoV-2 isolates require adaptation to use the mouse ACE-2 entry receptor and to productively infect the cells of the murine respiratory tract. Mouse-adapted strain of SARS-CoV-2 (MA10) causes infection, inflammation and pneumonia in BALB/c mice after intranasal inoculation **[11,13]**. MA10 has several mutations, including the N501Y mutation in the RBD of the spike protein compared to the Wuhan reference sequence, which also appears in the B.1.351 and B.1.1.7 variants **[6,7,11,23]**. These mutations in the RBD of the spike protein may have enhanced the binding affinity for the endogenous mouse ACE-2 receptor, thereby allowing the variants to replicate efficiently in mice. Indeed, studies have shown that SARS-CoV-2 variants containing N501Y and E484K mutations display substantially enhanced infection of mouse cells. Experimental studies using S glycoprotein and pseudo type viruses have shown that the B.1.1.7 and the B.1.351 variants bind to mouse ACE2 with higher affinity than the B.1 virus strain. In addition, B.1.351 virus has higher affinity for the mouse receptor than the B.1.1.7 [23-28]. The higher virus replication and inflammatory response triggered by the B.1.351 virus in mice agree with its higher affinity for the mouse receptor than the B.1.1.7 virus.

It has been suggested that a cytokine storm is involved in the pathogenesis of severe COVID-19 cases **[20,21]**. The levels of many cytokines and chemokines have been found to be increased after SARS-CoV-2 infection in humans. In the present study, we show that B.1.351 and B.1.1.7 infection induced significantly higher levels of cytokines and chemokine expression in the lungs. Inflammatory response observed in the B.1.351-infected group was higher compared to other groups. These findings are consistent with previous studies in humans, which establish a consistent link between the mutations exhibited by the B.1.351 lineage and a greater potential for infectivity and immune escape [4,6,24]. More studies are needed to characterize the pathological consequences of infection with these variants in different mouse strains and mice with co-morbid conditions. It is possible that more severe condition could be observed in mice with co-morbid conditions such as old age, diabetes and hypertension. The ability of clinical SARS-CoV-2 isolates to replicate and induce inflammation in wild-type mice will facilitate studies to evaluate therapeutic interventions and pathogenesis studies. These data indicate the possibility of adaptation to new animal species by emerging SARS-CoV-2 variants.

## Acknowledgments

We thank members of the GSU High Containment Core and the Department for Animal Research for assistance with the experiments.

